# Using dynamic oral dosing of rifapentine and rifabutin to simulate exposure profiles of long-acting formulations in a mouse model of tuberculosis preventive therapy

**DOI:** 10.1101/2023.04.12.536604

**Authors:** Yong S. Chang, Si-Yang Li, Henry Pertinez, Fabrice Betoudji, Jin Lee, Steven P. Rannard, Andrew Owen, Eric L. Nuermberger, Nicole C. Ammerman

**Author notes:** These authors contributed equally to this work (Yong S. Chang, Si-Yang Li). Corresponding author: Nicole C. Ammerman.

## Abstract

Administration of tuberculosis preventive therapy (TPT) to individuals with latent tuberculosis infection is an important facet of global tuberculosis control. The use of long-acting injectable (LAI) drug formulations may simplify and shorten regimens for this indication. Rifapentine and rifabutin have anti-tuberculosis activity and physiochemical properties suitable for LAI formulation, but there are limited data available for determining the target exposure profiles required for efficacy in TPT regimens. The objective of this study was to determine exposure-activity profiles of rifapentine and rifabutin to inform development of LAI formulations for TPT. We utilized a validated paucibacillary mouse model of TPT in combination with dynamic oral dosing of both drugs to simulate and understand exposure-activity relationships to inform posology for future LAI formulations. This work identified several LAI-like exposure profiles of rifapentine and rifabutin that, if achieved by LAI formulations, could be efficacious as TPT regimens and thus can serve as experimentally-determined targets for novel LAI formulations of these drugs. We present novel methodology to understand the exposure-response relationship and inform the value proposition for investment in development of LAI formulations that has utility beyond latent tuberculosis infection.

## INTRODUCTION

Tuberculosis (TB) is a major cause of morbidity and mortality worldwide. Control of TB is complicated for many reasons, including that the causative agent, *Mycobacterium tuberculosis*, can establish latency, referred to as latent TB infection (LTBI), which can persist throughout a lifetime. Thus, people with LTBI are at risk for bacterial reactivation and development of active TB disease, which is associated with individual morbidity and mortality, but also facilitates community transmission of *M. tuberculosis*. Fortunately, treatment for LTBI exists, termed TB preventive therapy (TPT) because it prevents the transition from latent infection to active disease. Up to one quarter of the global population are estimated to have LTBI, and appropriate administration of TPT is an essential component of the World Health Organization (WHO) End TB Strategy (1, 2).

Currently, there are five WHO-recommended TPT regimens for treatment of LTBI: 6-9 months of daily isoniazid, 4 months of daily rifampin, 3 months of daily isoniazid plus rifampin, 3 months of weekly isoniazid plus rifapentine, and one month of daily isoniazid plus rifapentine (1HP) (3). Although highly efficacious when treatment is completed, TPT is woefully underutilized. Among people with LTBI, it is estimated that only about 31% start TPT and 19% complete treatment (4). Unsurprisingly, treatment completion rates are inversely correlated with the duration of the TPT regimen, with the shorter regimens associated with the highest completion rates (3). Therefore, further shortening of TPT regimens could improve treatment completion rates. If development can be successfully accomplished, long-acting injectable (LAI) TPT offers the opportunity for a single administration at the point of care, ensuring adherence and treatment completion and simplifying programmatic roll-out.

Interest in the development of LAI formulations for treatment of LTBI has grown in recent years. The suitability of drugs for formulation as an LAI depends on certain physiochemical and pharmacokinetic (PK) properties, including low water solubility to prevent rapid release of the drug, low systemic clearance to enable sufficient concentrations to be achieved, and high potency to ensure target exposures are more easily reached and can be maintained for a protracted period of time. Swindells *et al.* recently summarized these properties among existing anti-TB drugs and found that for drugs used in the currently recommended TPT regimens, isoniazid and rifampin do not have properties amenable to LAI formulation while rifapentine is a promising candidate (5). In addition, Rajoli *et al.* used physiologically-based PK (PBPK) modeling to simulate potential LAI administration strategies for several anti-TB drugs, including rifapentine (6). This work identified a rifapentine exposure profile predicted to be both achievable with an LAI formulation and efficacious against LTBI. The efficacy prediction was based on maintaining plasma rifapentine levels above a specified target concentration of 0.18 μg/mL. This target concentration was defined as three times higher than a previously published minimum inhibitory concentration (MIC) of rifapentine for *M. tuberculosis*, but requires additional empirical validation (6, 7).

The physiochemical properties of another rifamycin antibiotic, rifabutin, appear even more favorable than rifapentine for LAI formulation (5, 8). Although only currently indicated for prevention and treatment of nontuberculous mycobacterial disease (9, 10), rifabutin has demonstrated potent anti-TB activity in preclinical models, including preclinical models of TPT (11-15), and has been used off-label for prevention and treatment of TB (reviewed in (16)). The suitability of rifabutin as a long-acting formulation is highlighted in a recent publication by Kim *et al.* describing the development of long-acting biodegradable *in situ* forming implants of rifabutin (13).

Dosing of one such formulation maintained plasma rifabutin concentrations above a target of 0.064 μg/mL for up to 18 weeks post-injection and exhibited robust activity in preclinical mouse models of TB treatment and infection prevention (*i.e*., pre-exposure prophylaxis). This target concentration was the epidemiological cut-off (ECOFF) from a previously reported MIC distribution of rifabutin against wild-type strains of *M. tuberculosis* (17, 18). However, because there were no recoverable colony forming units (CFU), the relationship between rifabutin plasma concentrations and activity could not be more precisely defined.

During development and preclinical evaluation of LAI formulations, selection of appropriate drug exposure profiles is critical for gauging therapeutic potential. However, there is a paucity of data regarding drug exposures required for efficacy in TPT, especially when considering the unique concentration-time profiles associated with a highly intermittent or single-shot LAI-based therapy compared to repeated daily or even weekly oral dosing. The objective of this project was to generate *in vivo* PK and pharmacodynamic (PD) data for rifapentine and rifabutin to inform LAI formulation development of these drugs for use in TPT. We designed orally-dosed regimens of rifapentine and rifabutin to produce specific plasma exposure profiles in BALB/c mice, and we evaluated the bactericidal activity of these regimens in a validated paucibacillary mouse model of TPT, a model that successfully predicted the efficacy of the 1HP TPT regimen in humans (19-21) and is the basis for translational PK/PD modeling of rifapentine activity in TPT regimens (22). The series of PK and PK/PD experiments described here provides a rich, experimentally-determined evidence base for selection of target exposure profiles which can help guide the development of LAI formulations of rifapentine and rifabutin for LTBI.

## RESULTS

### *In vitro* activity of rifamycins against *M. tuberculosis* H37Rv

We first determined the MIC and minimum bactericidal concentration (MBC) of rifapentine and rifabutin for our lab stock of *M. tuberculosis* H37Rv, with rifampin included as a comparator. As expected, the MIC rank order was rifampin (0.125-0.25 μg/mL) > rifapentine (0.0625 μg/mL) > rifabutin (0.0156 μg/mL). The observed MIC for rifapentine aligned with the MIC used in the modeling work by Rajoli *et al*. (6). For each rifamycin, the visual MIC and the MBC were the same concentration, and based on CFU counts, the concentration that inhibited bacterial growth was at least one 2-fold dilution lower than the visual MIC (**Table S1**).

### Plasma exposure profiles of low-dose oral rifapentine and rifabutin in uninfected BALB/c mice

Previous modeling work by Rajoli *et al.* utilized rifapentine at 0.18 μg/mL, or 3 times the MIC (3xMIC), as a target trough plasma concentration against which to measure predicted LAI rifapentine exposures (6). Thus, we were interested to evaluate *in vivo* exposure-activity relationships at mouse plasma concentrations around 3xMIC for both rifapentine and rifabutin. For both rifamycins, available oral PK data in mice was based on doses that yielded much higher plasma drug concentrations (11, 20, 23-25). In order to establish dose-exposure profiles within a relatively lower dosing range, we conducted pilot PK studies with rifapentine and rifabutin in uninfected BALB/c mice. The PK parameters of both drugs in female mice are presented in **Table 1** with PK profile data for both drugs being adequately described with a 1-compartment PK disposition model with 1^st^ order absorption input. For rifapentine, the maximum observed plasma concentration (C_max_) and the area under the plasma concentration versus time curve from 0-24 hours post-dose (AUC_0-24_) exhibited approximate dose linearity (**Fig. S1A**), with both parameters about 1.5-1.6 times higher on Day 10 compared to Day 0. While apparent clearance and plasma half-life remained relatively constant across the dose levels, the apparent volume of distribution and the absorption rate constant (Ka) decreased with increasing rifapentine dose. For rifabutin, the C_max_ and AUC_0-24_ also exhibited approximate dose linearity (**Fig. S1B**), and these parameters remained relatively stable from Day 0 to Day 10 of dosing (**Table 1**). Compared to rifapentine, rifabutin apparent clearance was much higher for all doses, and the plasma half-life was much shorter; apparent volume of distribution and Ka were also much higher, with the volume remaining relatively stable and Ka nominally decreasing with increasing dose but indicating very rapid absorption at all dose levels. For both rifapentine and rifabutin, no significant differences were observed for any of the PK parameters determined in male mice (**Table S2**). All individual mouse PK data are provided in **Supplementary Data File S1**. The available data from this study were used to generate a PK model to guide regimen design in the subsequent PK/PD study.

**Table 1.** PK parameters^a^ associated with oral dosing of rifapentine and rifabutin in uninfected female BALB/c mice. All individual mouse PK data are provided in **Data File S1**.

| Drug | Dose<sup>b</sup><br>(mg/kg) | Sampling<br>day<sup>c</sup> | T<sub>max</sub><br>(h post dose) | C<sub>max</sub><br>(µg/mL) | AUC<sub>0-24h</sub><br>(µg.h/mL) | CL/F<br>(L/h/kg)<br>[% RSE] | V/F<br>(L/kg)<br>[% RSE] | Ka<br>(h<sup>-1</sup>)<br>[% RSE] | t<sub>½</sub><br>(h) |
| --- | --- | --- | --- | --- | --- | --- | --- | --- | --- |
| Rifapentine | 0.1 | 0 | 2 | 0.130 | 2.5 | 0.024<br>[3.9] | 0.743<br>[5.3] | 2.57<br>[21.0] | 21.5 |
|  |  | 10 | 1 | 0.199 | 3.6 |  |  |  |  |
|  | 0.3 | 0 | 2 | 0.386 | 6.4 | 0.029<br>[3.0] | 0.736<br>[4.7] | 1.75<br>[13.5] | 17.6 |
|  |  | 10 | 2 | 0.644 | 9.6 |  |  |  |  |
|  | 1 | 0 | 2 | 1.58 | 23.9 | 0.021<br>[4.7] | 0.643<br>[6.9] | 1.64<br>[19.4] | 21.2 |
|  |  | 10 | 1 | 2.37 | 38.7 |  |  |  |  |
|  | 3 | 0 | 2 | 6.69 | 105.7 | 0.016<br>[4.4] | 0.418<br>[7.0] | 1.33<br>[17.4] | 18.1 |
|  |  | 10 | 2 | 11.9 | 166.6 |  |  |  |  |
| Rifabutin | 0.3 | 0 | 0.5 | 0.053 | 0.4 | 0.674<br>[5.6] | 7.36<br>[7.9] | 49.10<br>[0.0] | 7.6 |
|  |  | 10 | 1 | 0.041 | 0.3 |  |  |  |  |
|  | 1 | 0 | 1 | 0.140 | 1.0 | 1.04<br>[4.7] | 7.49<br>[6.3] | 16.40<br>[0.0] | 5.0 |
|  |  | 10 | 0.5 | 0.105 | 0.8 |  |  |  |  |
|  | 3 | 0 | 1 | 0.325 | 2.7 | 1.17<br>[4.7] | 8.67<br>[6.1] | 8.48<br>[212.9] | 5.1 |
|  |  | 10 | 0.5 | 0.348 | 2.4 |  |  |  |  |

<sup>a</sup>T<sub>max</sub>, time point of maximum observed plasma concentration; C<sub>max</sub>, maximum observed plasma concentration; AUC<sub>0-24h</sub>, area under the plasma concentration versus time curve from 0-24 hours post-dose; CL/F, apparent clearance; V/F, apparent volume of distribution; Ka, absorption rate constant; t<sub>½</sub>, plasma half-life; % RSE, percent relative standard error.
<sup>b</sup>Drugs were administered once daily, five days per week (Monday-Friday).
<sup>c</sup>Day 0 indicates the first day of drug administration (*i.e.*, the first dose).

### First PK/PD study of rifapentine and rifabutin at steady, LAI-relevant plasma concentrations in the paucibacillary mouse model of TPT

To understand exposure-activity relationships of rifapentine and rifabutin, we conducted a PK/PD study to evaluate the *in vivo* bactericidal activity when drug exposures are maintained at steady, LAI-relevant plasma concentrations. For rifapentine, we selected four target trough plasma concentrations of interest for LAI formulations: 0.18 μg/mL (3xMIC); 0.6 μg/mL (the trough plasma concentration predicted by Rajoli *et al.* four weeks after a single 750 mg intramuscular injection of a simulated rifapentine LAI with a nominal first order release rate constant from the injection site of 0.0015 h^-1^ (6)); 3.5 μg/mL (the reported average plasma concentration of rifapentine in humans taking the 3-month, once-weekly, oral high-dose isoniazid and rifapentine TPT regimen (26)); and 2 μg/mL (a concentration selected to fill the exposure gap between 0.6 and 3.5 μg/mL). For rifabutin, we selected two target trough plasma concentrations of interest, namely 0.045 μg/mL (3xMIC) and 0.15 μg/mL (the reported average plasma concentration in people receiving an oral, once daily 150 mg dose (27, 28)).

After determining the target plasma concentrations of interest, we used our PK model to design oral dosing schemes, based on twice daily dosing, that were predicted to maintain relatively stable mouse plasma trough concentrations above each pre-defined target concentration (**Fig. S2A-F**). Because the difference in predicted peak-to-trough plasma concentrations between oral doses was relatively large for rifabutin, we designed two additional dosing schemes to maintain average plasma concentrations at the pre-defined targets of 0.045 and 0.15 μg/mL (**Fig. S2G-H**). The dosing schemes predicted to achieve each plasma target exposure are summarized in **Table 2**. The PK and bactericidal activity of these eight regimens were then evaluated in the paucibacillary mouse model of TPT, with once daily oral rifampin included as a rifamycin positive control regimen (see experiment scheme in **Table S3**).

**Table 2.** Regimens for the first PK/PD study. The full experiment scheme is presented in Table S3.

| Regimen description<sup>a</sup> | Oral dosing scheme<sup>b</sup> |
| --- | --- |
| **Control regimens** |  |
| Negative control | Untreated |
| Positive control | Rifampin 10 mg/kg QD |
| **Rifapentine (RPT) test regimens** |  |
| RPT target: C<sub>trough</sub> 0.18 µg/mL | RPT 0.075 mg/kg BID with 0.2 mg/kg loading dose |
| RPT target: C<sub>trough</sub> 0.6 µg/mL | RPT 0.25 mg/kg BID with 0.625 mg/kg loading dose |
| RPT target: C<sub>trough</sub> 2 µg/mL | RPT 0.75 mg/kg BID with 1.875 mg/kg loading dose |
| RPT target: C<sub>trough</sub> 3.5 µg/mL | RPT 1.25 mg/kg BID with 3.125 mg/kg loading dose |
| **Rifabutin (RFB) test regimens** |  |
| RFB target: C<sub>ave</sub> 0.045 µg/mL | RFB 0.75 mg/kg BID |
| RFB target: C<sub>trough</sub> 0.045 µg/mL | RFB 1.5 mg/kg BID |
| RFB target: C<sub>ave</sub> 0.15 µg/mL | RFB 2.5 mg/kg BID |
| RFB target: C<sub>trough</sub> 0.15 µg/mL | RFB 5.5 mg/kg BID |

<sup>a</sup>C<sub>trough</sub> indicates that the regimen was designed to maintain trough plasma concentrations at this target.
C<sub>ave</sub> indicates that the regimen was designed to maintain average plasma concentrations at this target.
<sup>b</sup>QD indicates once daily dosing; BID indicates twice daily dosing. All rifapentine regimens included a single loading dose administered as the first dose on Day 0 (the first day of treatment). All regimens were administered 7 days per week.

For rifapentine, the observed trough plasma concentrations and profiles in general, agreed reasonably with PK model simulations based on the pilot PK in uninfected mice, with the greatest discrepancy under the regimen with the lowest target concentration of 0.18 μg/mL (**Table 3**; **Fig. S3**). Observed trough plasma concentrations under these regimens aligned well with the target at the lower target trough concentrations of 0.18 and 0.6 μg/mL, (**Fig. S3A-B**), but exceeded the targets more at the higher targets of 2 and 3.5 μg/mL (**Figs S3C-D**). This was in part due to the unavailability (at the time of regimen design) of the Day 10 pilot PK exposure data for inclusion in the PK model fitting and parameter estimation. Had these data been available at the time of regimen design, somewhat lower twice-daily maintenance doses for the higher targets of 2 and 3.5 μg/mL would have been chosen. The rifapentine dosing schemes predicted to achieve trough plasma concentrations of 0.18 and 0.6 μg/mL (with data-fitted trough plasma concentrations of 0.17 and 0.54 μg/mL, respectively) did not exhibit any bactericidal activity during the three weeks of treatment (**Fig. 1**; **Table S4**). However, the dosing schemes predicted to achieve the higher two plasma trough targets of 2 and 3.5 μg/mL (with data-fitted trough plasma concentrations of 2.6 and 4.9 μg/mL, respectively) were associated with significant bactericidal activity, with *M. tuberculosis* killing equivalent to that of the rifampin positive control regimen. Rifapentine activity was dose-dependent, and this relationship was also reflected in the modeled bacterial elimination rate constant (k_net_) (**Table 3**; **Fig. S4A**). The relationship between k_net_ and observed drug exposures was similar (**Fig. S4B-C**).

**Fig. 1.**
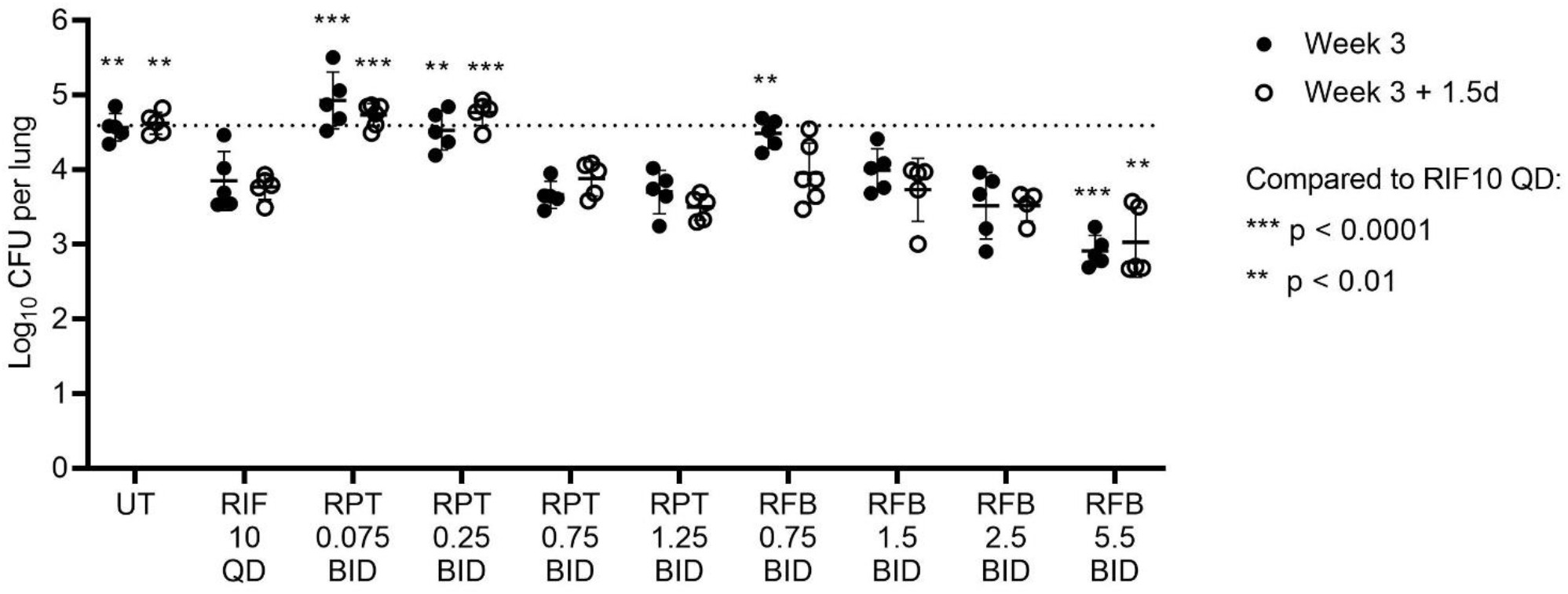
Bactericidal activity in the first PK/PD study. The original study scheme included 6 weeks of treatment, with bactericidal activity assessed at Week 3 and Week 6. However, the COVID-19-related university shut-down necessitated ending this study 1.5 days after the Week 3 time point. UT, untreated; RIF, rifampin; RPT, rifapentine; RFB, rifabutin; QD, once daily; BID, twice daily. Drug dose in mg/kg and frequency of administration are given below each drug abbreviation; see regimen descriptions in **Table 2**. Each data point represents and individual mouse, and lines/error bars represent the mean and standard deviation. The dotted horizontal line indicates the average lung CFU count at the start of treatment (Day 0). At each time point, lung CFU counts for all treatment groups were compared against lung CFU counts in the RIF 10 QD positive control group using 2-way ANOVA with Dunnett’s multiple comparisons test. Individual mouse CFU data and statistical analysis are provided in **Data File S2**.

**Table 3.**
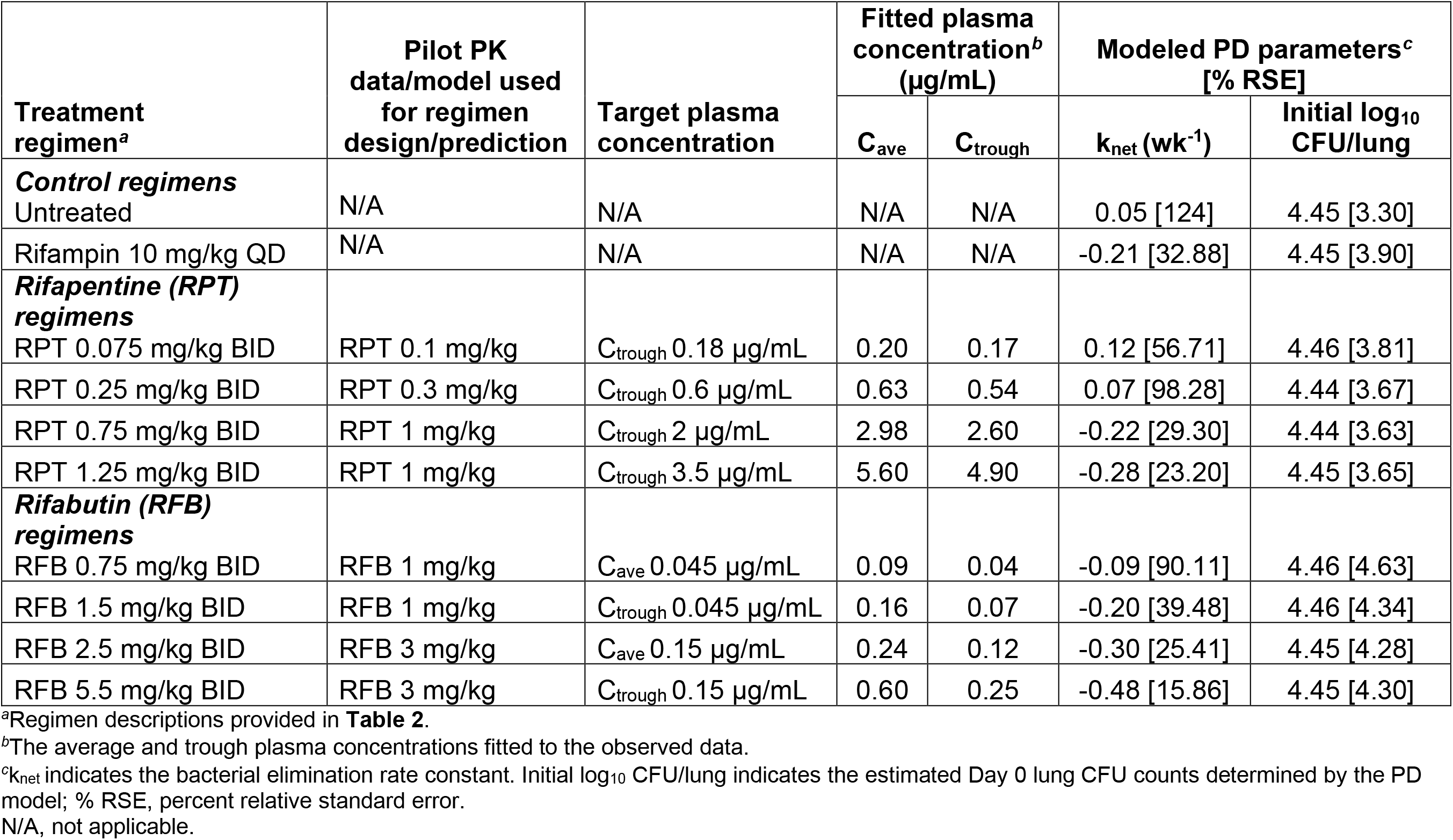
PK/PD parameters for rifapentine and rifabutin in the first PK/PD study. All individual mouse CFU and PK data are presented in **Data File S2**.

For rifabutin, the observed plasma concentrations across all regimens were slightly higher than concentrations predicted from the pilot PK model but otherwise generally in good agreement (**Table 3**; **Fig. S5**). The dosing scheme predicted to achieve a trough plasma target concentration of 0.045 μg/mL (data-fitted trough concentration 0.07 μg/mL) had limited bactericidal activity (**Fig. 1**; **Table S4**). Other rifabutin dosing schemes were associated with further dose-dependent bactericidal activity. The dosing schemes predicted to achieve either a target average plasma concentration of

0.045 μg/mL or a trough concentration of 0.15 μg/mL (with data-fitted average and trough plasma concentrations of 0.09 and 0.25 μg/mL, respectively) exhibited bacterial killing equivalent to the rifampin control regimen, and the dosing scheme predicted to achieve an average plasma target of 0.15 (data-fitted average concentration 0.24 μg/mL) was associated with significantly greater *M. tuberculosis* killing than the rifampin control. Across the regimens, there were strong dose-dependent and exposure-dependent effects on bactericidal activity (**Table 3**; **Fig. S4D-F**). For all regimens in this study, the PD model estimated that the bacterial lung burden at the start of treatment was 4.44-4.46 log_10_ CFU/lung, which aligned with the observed Day 0 mean lung burden of 4.47 log_10_ CFU/lung.

### Second PK/PD study with dynamic oral dosing of rifapentine and rifabutin to simulate possible LAI exposure profiles in the paucibacillary mouse model of TPT

Our hitherto collected PK/PD data, as well as predicted human PBPK parameters from previous modeling work (6), were used to generate simulated LAI-like exposure profiles of rifapentine and rifabutin expected to have bactericidal activity that could be achieved using oral dosing with a dynamic, tapered dose regimen design. These profiles simulated the initial increase and then relatively slow decline of plasma drug concentrations that could occur following an intramuscular injection of an LAI formulation. The profiles were designed to maintain trough plasma concentrations above pre-defined target concentrations (0.6, 2, and 3.5 μg/mL for rifapentine; and 0.045 and 0.15 μg/mL for rifabutin) by 4 weeks after each simulated LAI dose. For the rifapentine target trough concentrations of 0.6 and 2 μg/mL, exposure profiles were also generated to simulate two LAI doses given 4 weeks apart. To test the bactericidal activity associated with each exposure profile in the paucibacillary mouse model, orally-dosed regimens were designed to achieve each profile. To capture the dynamic nature of the simulated LAI exposures over time, the oral rifamycin doses changed every 4 days. The desired rifapentine and rifabutin simulated LAI exposure profiles and the simulated plasma concentrations associated with each oral regimen are shown in **Fig. S6** (rifapentine) and **Fig. S7A-B** (rifabutin). The regimens are summarized in **Table 4**, with detailed descriptions of the dosing provided in **Table S5**. The PK and bactericidal activity associated with each simulated LAI regimen was then evaluated in the paucibacillary mouse model (see the experimental scheme in **Table S6**).

**Table 4.** Regimens for the second PK/PD study. The full experiment scheme is presented in Table S6.

| Simulated LAI treatment regimen<sup>a</sup> | Oral dosing regimen<sup>b</sup> for LAI simulation during the indicated treatment period: |  |
| --- | --- | --- |
|  | Weeks 1-4 | Weeks 5-8 |
| Negative control | Untreated | Untreated |
| Positive control: 1HP | INH 10 mg/kg QD + RPT 10 mg/kg QD |  |
| RPT LAI target 0.6 µg/mL, 1 dose | RPT 0.75 mg/kg BID to 0.275 mg/kg BID | RPT 0.24 mg/kg BID to 0.1 mg/kg BID |
| RPT LAI target 0.6 µg/mL, 2 doses | RPT 0.75 mg/kg BID to 0.275 mg/kg BID | RPT 1 mg/kg BID to 0.375 mg/kg BID |
| RPT LAI target 2 µg/mL, 1 dose | RPT 1.6 mg/kg BID to 0.6 mg/kg BID | RPT 0.5 mg/kg BID to 0.2 mg/kg BID |
| RPT LAI target 2 µg/mL, 2 doses | RPT 1.6 mg/kg BID to 0.6 mg/kg BID | RPT 2.1 mg/kg BID to 0.8 mg/kg BID |
| RPT LAI target 3.5 µg/mL, 1 dose | RPT 2.4 mg/kg BID to 0.875 mg/kg BID | RPT 0.8 mg/kg BID to 0.325 mg/kg BID |
| RFB LAI target 0.045 µg/mL, 1 dose | RFB 1.5 mg/kg BID to 0.6 mg/kg BID | RFB 0.5 mg/kg BID to 0.2 mg/kg BID |
| RFB LAI target 0.15 µg/mL, 1 dose | RFB 5 mg/kg BID to 2.2 mg/kg BID | RFB 2 mg/kg BID to 0.8 mg/kg BID |

<sup>a</sup>The positive control, 1HP, indicates 1 month (4 weeks) of isoniazid (INH) and rifapentine (RPT); RFB, rifabutin. Simulated long-acting injectable (LAI) regimens were designed to maintain trough plasma concentrations above the indicated target for 4 weeks per simulated LAI dose.
<sup>b</sup>QD indicates once daily dosing; BID indicates twice daily dosing; all regimens were administered 7 days per week. For simulated LAI regimens, the dosing changed every 4 days. For regimens simulating one LAI dose, the dosing decreased continuously during all 8 weeks of treatment. For regimens simulating two LAI doses, the dosing decreased during weeks 1-4, but then increased again at week 5, following by a continuous decrease through week 8. The dosing range provided in the table represents the starting and ending doses for weeks 1-4 and 5-8 of treatment. See **Table S5** for a full description of the dosing changes for each simulated LAI regimen.

For the simulated LAI regimens of rifapentine, the exposure profiles fitted from the observed data aligned well with the predicted exposures (**Fig. 2**; **Table 5**), although data-fitted exposures trended slightly lower than predicted exposures both with increasing rifapentine dose and increasing duration of administration. This is in keeping with known autoinduction of clearance by rifamycins on this timescale of administration which was not captured in the shorter duration pilot PK data and model from which these regimens were designed. In fitting PK models to the study 2 plasma PK exposure profiles, the estimated “induction factor” for time dependent, % daily (cumulative) increase in clearance was approximately +1% per dosing day for both rifapentine and rifabutin under all regimens (**Table 5**) leading to ∼1.7x higher clearance by day 52 of the study compared to day 1. Our goal was to maintain trough plasma levels above the pre-determined target concentrations for 4 weeks following each simulated LAI dose, and this was achieved or nearly achieved for all regimens. All simulated rifapentine LAI regimens exhibited bactericidal activity in the mouse lungs, with clear dose-dependent killing effects (**Fig. 3A**; **Table S7**). A single simulated LAI dose for all target concentrations was significantly bactericidal compared to the untreated mice at Week 4 (p < 0.001 for all regimens), and the regimen with a plasma trough concentration target of 3.5 μg/mL had equivalent killing as the 1HP control regimen at Weeks 2 and 4, and a modest additional killing effect between Weeks 4 and 8. Additionally, two doses of the regimen with a plasma trough concentration target of 2 μg/mL achieved the same killing effect at Week 8 as the 1HP control regimen did at Week 4.

**Fig. 2.**
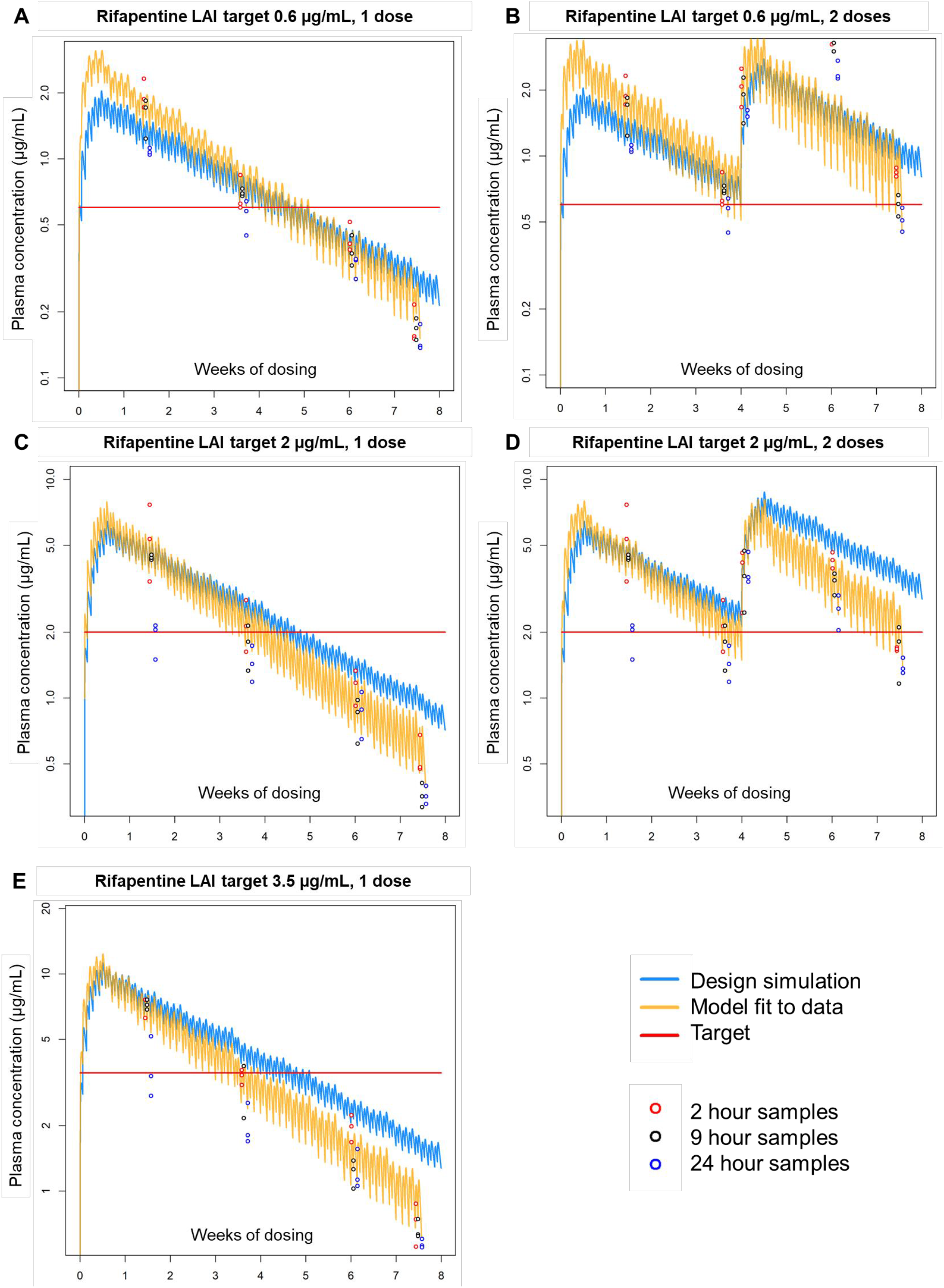
Simulated, observed, and modeled rifapentine PK data from the second PK/PD study. PK data for simulated LAI regimens with targets of 0.6, 2, and 3.5 μg/mL are shown in panels A-B, C-D, and E, respectively. In each graph, the simulated exposure profile based on the regimen design is shown in blue, the target plasma concentrations are shown by the red line, the observed data points are indicated by the open circles, and the modeled exposure profiles fitted to the observed data are shown in yellow. For the regimen with a rifapentine LAI target of 0.6 μg/mL with two simulated doses (panel B), the 9-hour PK data were omitted from the model fitting. Individual mouse PK data are provided in **Data File S3**.

**Fig. 3.**
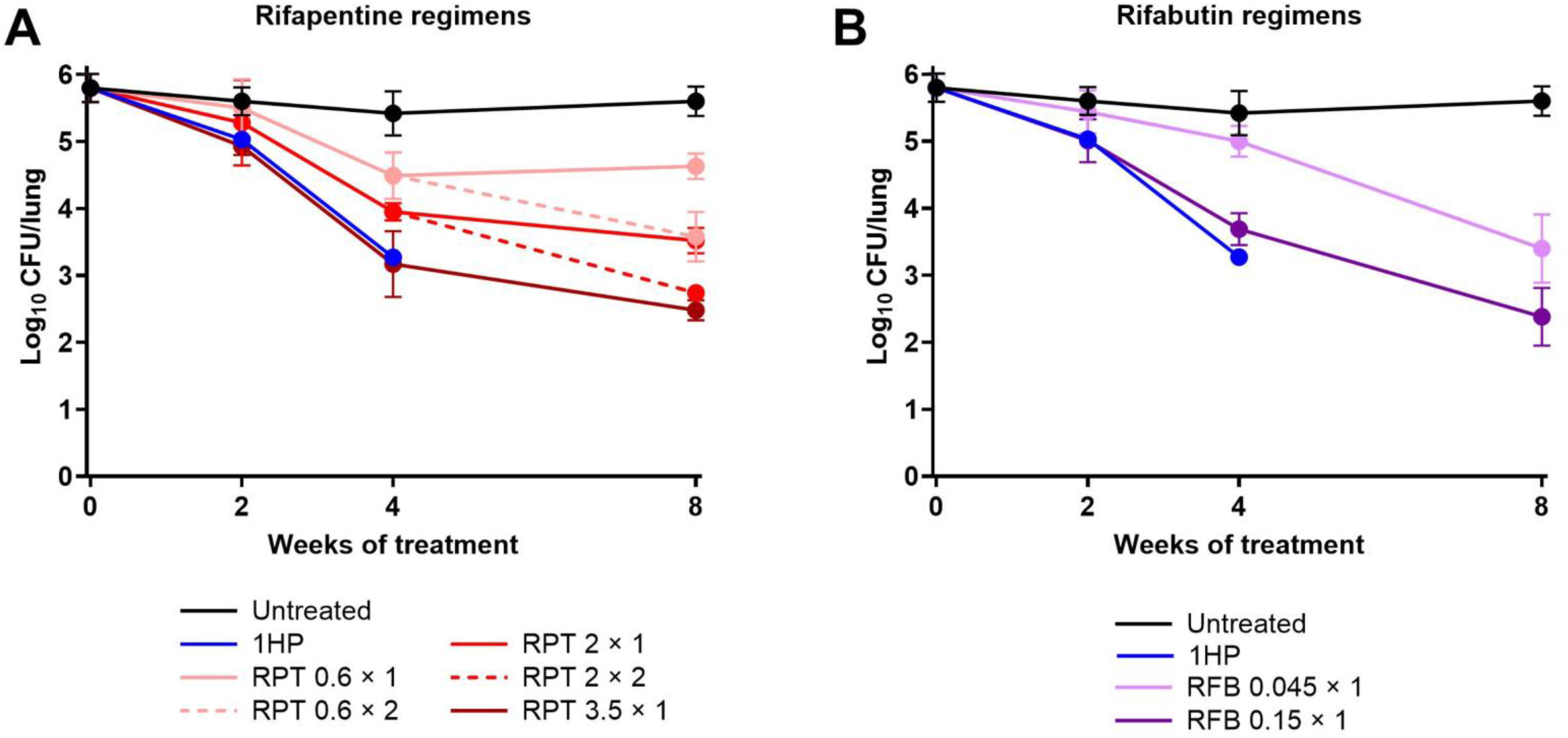
Bactericidal activity of simulated LAI regimens with rifapentine (A) and rifabutin (B) in the second PK/PD study. 1HP, control regimen (once daily isoniazid and rifapentine); RPT, rifapentine; RFB, rifabutin. For rifapentine and rifabutin simulated LAI regimens, the target plasma concentration in μg/mL and the number of simulated doses (× 1 or × 2) are given after each drug abbreviation; see regimen descriptions in **Table 4**. Data points represent the mean, and error bars represent the standard deviation. Individual mouse CFU data and statistical analysis are provided in **Data File S3**.

**Table 5.** PK and dosing information for second PK/PD study, with simulated rifapentine and rifabutin LAI regimens.

| Simulated LAI treatment regimen<sup>a</sup> | PK data/model used for regimen design/prediction<sup>b</sup> | Total dose (mg/kg) | Trough plasma concentration<sup>c</sup> (µg/mL) |  | Cumulative AUC (µg·h/mL)<sup>d</sup> |  | CL/F (L/h/kg) [% RSE] | V/F (L/kg) [% RSE] | Ka (h<sup>-1</sup>) [% RSE] | Ind_Factor (%) [%RSE] |
| --- | --- | --- | --- | --- | --- | --- | --- | --- | --- | --- |
|  |  |  | Day 28 | Day 52 | Predicted | Fitted |  |  |  |  |
| RPT LAI target 0.6 µg/mL, 1 dose | RPT 0.25 mg/kg | 35.4 | 0.60 | 0.15 | 1078 | 1320 | 0.02 [9.6] | 0.37 [122.0] | 0.23 [128.9] | 1.1 [17.8] |
| RPT LAI target 0.6 µg/mL, 2 doses | RPT 0.25 mg/kg | 61.8 | 0.52 | 0.45 | 1866 | 2093 | 0.02 [4.7] | 0.35 [16.8] | 0.42 [23.3] | 1.1 [11.9] |
| RPT LAI target 2 µg/mL, 1 dose | RPT 0.75 mg/kg | 74.0 | 1.58 | 0.38 | 3558 | 3251 | 0.02 [9.4] | 0.64 [29.9] | 5.09 [0.004] | 1.1 [23.4] |
| RPT LAI target 2 µg/mL, 2 doses | RPT 0.75 mg/kg | 129.2 | 1.52 | 1.37 | 6143 | 5074 | 0.02 [7.8] | 0.48 [25.5] | 0.62 [46.3] | 1.2 [19.5] |
| RPT LAI target 3.5 µg/mL, 1 dose | RPT 1.25 mg/kg | 113.8 | 2.32 | 0.59 | 6182 | 5217 | 0.02 [12.6] | 0.45 [70.4] | 0.57 [101.4] | 1.1 [24.7] |
| RFB LAI target 0.045 µg/mL, 1 dose | RFB 2.5 mg/kg | 70.4 | 0.03 | 0.01 | 91 | Not calculated<sup>e</sup> | Not fitted<sup>e</sup> | Not fitted<sup>e</sup> | Not fitted<sup>e</sup> | Not fitted<sup>e</sup> |
| RFB LAI target 0.15 µg/mL, 1 dose | RFB 2.5 mg/kg | 254.8 | 0.06 | 0.02 | 274 | 200 | 1.06 [15.6] | 13.23 [19.8] | 10.03 [0.5] | 0.9 [46.4] |

<sup>a</sup>RPT, rifapentine; RFB, rifabutin. Simulated long-acting injectable (LAI) regimens were designed to maintain trough plasma concentrations above the indicated target for 4 weeks (28 days) per simulated LAI dose. See **Table 4** for a description of the regimens.
<sup>b</sup>Twice daily (BID) dosing regimen from PK/PD Study 1
<sup>c</sup>The trough plasma concentration (modeled from fitting to the data) achieved at the stated time into the simulated LAI dose profile (for comparison against the stated target concentration). For RFB LAI target 0.045 µg/mL 1 dose regimen, predicted values reported (see note e).
<sup>d</sup>Area under the concentration-time curve (AUC) that was simulated prior to the experiment (see blue lines in **Fig. 2** and **Fig. S7**) and fitted from the observed data (see yellow lines in **Fig. 2** and **Fig. S7**).
<sup>e</sup>Cumulative AUC was not calculated, and model fitting parameter estimates are not reported due to poor model fit due to truncated PK data profile (see **Fig. S7**).
CL/F, apparent clearance on Day 1; V/F, apparent volume of distribution; Ka, absorption rate constant; Ind\_Factor, factor for percent cumulative increase in CL/F per dosing day; % RSE, percent relative standard error.

For the simulated LAI regimens of rifabutin, modeling of the drug exposures based on the observed data was limited because rifabutin was undetectable in many of the 9 h and 24 h plasma samples (**Fig. S7C-D**). However, the available observed data indicate that rifabutin plasma exposures aligned well with the predicted exposures for each regimen (**Table 5**). Both rifabutin LAI regimens were associated with significant bactericidal activity in the mouse model of TPT (**Fig. 3B; Table S7**). At Week 4, the simulated LAI regimen based on the Week 4 plasma trough concentration target of 0.15 μg/mL was as bactericidal as the 1HP control regimen at Weeks 2 and 4. This rifabutin regimen continued to exert bactericidal activity up to Week 8, resulting in a mean lung CFU count lower than that observed at the completion of the 1HP regimen. Despite having no significant bactericidal activity during the first 2 weeks of treatment, the simulated rifabutin LAI regimen based on the Week 4 plasma trough concentration target of 0.045 μg/mL exhibited potent bactericidal activity between Week 2 and Week 8 of treatment.

Due to the relatively low bacterial burden at the start of treatment, selection of rifamycin-resistant isolates was not expected in this study, and this expectation was confirmed. At Week 8, all samples were evaluated for rifamycin resistance by directly plating a portion of the lung homogenates on agar containing 1 μg/mL rifampin. No rifamycin resistance was detected in any sample (**Data File S3**).

## DISCUSSION

In this project, we used oral dosing of rifapentine and rifabutin in a validated paucibacillary mouse model of LTBI treatment to better understand the PK/PD relationships driving their activity and to gain insights into efficacious exposure profiles to help guide the development of LAI formulations of these drugs for TPT. For both rifamycins, we identified 4- and 8-week exposure profiles that had equivalent bactericidal activity to that of the 1HP control regimen in the mouse model. These data define exposure profiles of rifapentine and rifabutin that, if achieved by LAI formulations, could be as efficacious as existing TPT regimens. Thus, the presented data can serve as experimentally-determined targets for novel LAI formulations of these drugs.

A paucity of data exists to describe dose- or exposure-response relationships for TPT regimens against which to benchmark the desired PK profiles of novel LAI formulations. Furthermore, in the absence of an efficient, quantitative biomarker of clinical response to TPT, it is currently exceedingly challenging to define meaningful PK/PD relationships using clinical data. When Rajoli and colleagues used PBPK models to simulate LAI administration strategies of anti-TB drugs, they necessarily relied on *in vitro* characteristics to define target concentrations, such as 3xMIC for rifapentine, as is customary in early medicine development (6). Model-based simulations indicated that a 250 mg LAI dose of rifapentine would generate peak plasma concentrations around 0.5 μg/mL and would maintain rifapentine concentrations above the 3xMIC target concentration for at least 4 weeks post-injection. However, our data from the first PK/PD study indicate that maintaining rifapentine trough plasma concentrations *in vivo* between a trough of 0.17 μg/mL (around 3xMIC) and a peak of 0.22 μg/mL (**Fig. S3A**) or even between a trough of 0.54 μg/mL and a peak of 0.75 μg/mL (**Fig. S3B**) was not bactericidal in the paucibacillary mouse model of TPT (**Fig. 1**; **Table 3**). In contrast, dosing regimens maintaining rifapentine plasma concentrations of approximately 2.6-3.75 μg/mL (**Fig. S3C**) and 4.9-7 μg/mL (**Fig. S3D**) exerted increasing bactericidal activity. These findings are in line with prior evidence of a linear concentration-response between 2 and 10 μg/mL and little effect of concentrations below 1 μg/mL (16x MIC) in this model (22). Based on these data, an LAI target concentration of 3xMIC is not appropriate for rifapentine, and little bactericidal effect can be expected from concentrations below 16x MIC. However, in the case of rifabutin, maintaining a plasma concentration at between 0.07 μg/mL (4x MIC) and 0.25 μg/mL (16x MIC) was associated with significant bactericidal activity equivalent to that of the rifampin 10 mg/kg positive control regimen (**Fig. 1**; **Table 3**). Thus, despite both being rifamycins, basing the plasma target concentration on the same multiple of the MIC (which was also the MBC) did not accurately predict *in vivo* bactericidal activity. For rifapentine, our data suggest that cumulative AUC may align better with the observed bactericidal activity in mice (**Table 5**); we did not have sufficient data from the second PK/PD study to assess any relationship between rifabutin cumulative AUC and bactericidal activity. The possibility of an AUC target should therefore be investigated in future studies.

Our exposure-activity data for rifabutin align with that reported by Kim *et al.*, who found that a long-acting biodegradable *in situ* forming implant of rifabutin that maintained plasma concentrations above 0.064 μg/mL, a reported ECOFF concentration (17, 18) selected as a target for up to 18 weeks post-injection, exhibited potent anti-TB activity in mouse models (13). The formulation developed by Kim *et al.* is therefore especially promising given that our data suggest that far lower target concentrations of rifabutin may be sufficient for efficacy in TPT regimens. Interestingly, Rajoli *et al.* also used a reported ECOFF concentration of 1.6 μg/mL for bedaquiline as the desired target concentration against which to benchmark activity of simulated bedaquiline LAI doses based on PBPK modeling (6, 29). In this case, the maximum predicted exposure profile could not reach the target concentration, thus raising questions about the utility of an LAI formulation of bedaquiline for TPT. The *in vivo* evaluation of an actual bedaquiline LAI, an aqueous microsuspension developed by Janssen, demonstrated that a single 160 mg/kg injection in mice resulted in a plasma exposure profile that remained above the bedaquiline MIC (0.03 μg/mL) for up to 12 weeks post-injection but was indeed well below the ECOFF concentration of 1.6 μg/mL (30). However, this exposure profile was associated with good bactericidal activity, similar to that of rifampin monotherapy in the paucibacillary mouse model of TPT. Thus, taking together our findings along with those of previously published studies, it is clear that a standard, set target concentration such as 3xMIC or ECOFF, cannot be used across the board as exposure targets for LAI formulations, but rather target exposure profiles need to experimentally determined individually for each drug.

This raises the key question: how should target concentrations or exposure profiles of LAI formulations be determined for TPT? The development of LAI formulations is a long, difficult, and resource-intensive process. Therefore, knowledge of the necessary exposure profiles required for efficacy is invaluable to guide LAI development efforts. In this project, we present a novel approach to utilize oral dosing to simulate LAI exposure and determine *in vivo* exposure-activity relationships. However, the PK associated with oral dosing, even when dosed in a manner to try to maintain stable plasma concentrations or to mimic the extended rise and fall of a simulated LAI exposure profile, may not be directly comparable. The peaks and troughs associated with oral dosing may affect drug activity in a way that steady drug release from an LAI depot may not (and vice versa). Drug exposure may also be different in that the up- or down-regulation of gut and liver transporters can significantly impact the plasma concentration of orally-dosed drugs via both absorption/bioavailability and clearance mechanisms. For example, in this study, rifapentine levels decreased in a dose- and time-dependent manner compared to the simulated exposures (**Fig. 2**), consistent with this drug’s well-documented autoinduction of clearance in humans and in mice (22). Some of the PK issues specific to oral dosing could be mitigated by parenteral administration, but this would not eliminate the peaks and troughs associated with daily or twice daily dosing and would cause considerably more discomfort in the mice. Ultimately, only when an LAI formulation is available will an understanding of how well these oral dosing schemes align with exposure/activity relationships observed with actual LAI formulations emerge. Thus, the data from this current study are not providing definitive targets but rather can be used to guide LAI formulation development for rifapentine and rifabutin. Future studies with LAI formulations will be needed to confirm their validity.

Keeping in mind that oral dosing was used in this study, there are several interesting observations worth noting. First, the observed bactericidal activity of rifabutin in the second PK/PD study was strongest between Week 4 and Week 8 (**Fig. 3B**), when rifabutin plasma concentrations fell below the target concentrations and were even undetectable at many of the PK time points (**Fig. S7**). It is possible that plasma rifabutin concentrations remained above the MIC (0.0156 μg/mL), as this was below the lower limit of quantification (0.05 μg/mL). However, as only one simulated LAI dose was tested in the rifabutin regimens, the dosing steadily decreased over the 8 weeks of the study (**Table 4**), and therefore it is surprising that the most potent bactericidal activity occurred during the latter phase of rifabutin dosing. It should be noted that the lung tissue, *i.e.*, effect compartment, PK was not evaluated within our study and could provide a rational basis for these findings. Thus, these data add to the promise of rifabutin in the context of TPT, but also highlight the need for further PK/PD studies to better understand how this drug exerts its activity *in vivo*.

A second interesting observation was the nature of the bactericidal activity of rifapentine in the second PK/PD study (**Fig. 2A**). We tested three oral rifapentine regimens that simulated one LAI dose administered at Day 0. For each of these regimens, dose-dependent bactericidal activity was observed during the first 4 weeks, but between Week 4 and Week 8, bactericidal activity seemed to stall or end for all doses. Incorporation of a second simulated-LAI dose at Week 4 maintained bactericidal activity, also in a dose-dependent manner. However, when examining plasma rifapentine concentrations, the PK/PD relationships become less clear between Week 4 and Week 8. For example, in mice that received one simulated LAI dose designed to reach a trough of 2 μg/mL at Week 4, plasma rifapentine concentrations fell from around 2 μg/mL to around 0.5 μg/mL between Weeks 4 and 8 (**Fig. 2C**), and this was associated with stalled bactericidal activity (**Fig. 3A**). This same plasma exposure range was achieved in mice receiving one simulated LAI dose designed to reach a trough of 0.6 μg/mL at Week 4 (**Fig. 2A**), but in this case was associated with significant bactericidal activity. Thus, an important question is whether this pattern of bactericidal activity is inherent to the overall exposure profile and could be expected when dosing a real LAI formulation.

Overall, the use of dynamic oral dosing to mimic possible LAI exposures has revealed remarkable PK/PD relationships and promising exposure profiles for both rifapentine and rifabutin for treatment of LTBI. The data from this study support the development of LAI formulations of each of these rifamycins for use in TPT regimens, and if validated, this approach may help accelerate development of LAIs by providing evidence-based targets for preclinical studies in advance of or in parallel to initial formulation development.

## MATERIALS AND METHODS

All *in vitro* and *in vivo* experiments were conducted at the Johns Hopkins University School of Medicine. PK/PD analyses and modeling work were performed at the University of Liverpool.

### Mycobacterial strains

*M. bovis* rBCG30 (originally provided by Marcus A. Horwitz from the University of California – Los Angeles School of Medicine) (31) and *M. tuberculosis* H37Rv (originally purchased from American Type Culture Collection, strain ATCC 27294) were each mouse-passaged and frozen in aliquots.

### Media

The standard liquid growth medium for both mycobacterial strains was Middlebrook 7H9 broth supplemented with 10% (v/v) oleic acid-albumin-dextrose-catalase (OADC) enrichment, 0.5% (v/v) glycerol, and 0.1% (v/v) Tween 80. MIC/MBC assay medium was Middlebrook 7H9 broth supplemented with 10% (v/v) OADC and 0.5% (v/v) glycerol, without Tween 80. 7H11 agar supplemented with 10% (v/v) OADC and 0.5% (v/v) glycerol was the solid medium base for all samples. Lung homogenates and their cognate dilutions were cultured on 7H11 agar rendered selective for mycobacteria by the addition of 50 µg/mL carbenicillin, 10 µg/mL polymyxin B, 20 µg/mL trimethoprim, and 50 µg/mL cyclohexamide (32). Selective 7H11 agar was also supplemented with 0.4% (w/v) activated charcoal to detect and limit the effects of drug carryover in lung homogenates (33, 34). To differentiate *M. bovis* rBCG30 and *M. tuberculosis* H37Rv CFUs, selective 7H11 agar was further supplemented with either 40 µg/mL hygromycin B (selective for *M. bovis* rBCG30 but not *M. tuberculosis*) or 4 µg/mL 2-thiophenecarboxylic acid hydrazide (TCH, selective for *M. tuberculosis* but not *M. bovis*). Because TCH is adsorbed and rendered less active by activated charcoal (30), only hygromycin B (at 40 µg/mL) was added to charcoal-containing selective 7H11 agar. Difco Middlebrook 7H9 broth powder, Difco Mycobacteria 7H11 agar powder, and BBL Middlebrook OADC enrichment were obtained from Becton, Dickinson and Company. Glycerol and Tween 80 were obtained from Fisher Scientific, and activated charcoal was obtained from J. T. Baker. All selective drugs were obtained from Sigma-Aldrich/Millipore-Sigma.

### Drug sourcing, formulations, and administration

Rifampin and isoniazid powders were purchased from Millipore-Sigma. Rifapentine tablets (brand name Priftin®, manufactured by Sanofi) were purchased from the Johns Hopkins Hospital Pharmacy. Rifapentine and rifabutin powders were purchased from Biosynth. For MIC/MBC assays, rifampin, rifapentine (powder), and rifabutin were dissolved in dimethyl sulfoxide (DMSO). Rifampin was also dissolved in DMSO when added to agar for direct susceptibility testing of mouse lung homogenates. For administration to mice, rifampin, rifapentine (tablets), and rifabutin were formulated in 0.05% (w/v) agarose in distilled water; isoniazid was dissolved in distilled water. All drugs were formulated to deliver the indicated oral dose in a total volume of 0.2 mL, administered by gavage. Dosing by mg/kg was based on an average mouse body mass of 20 g.

### MIC/MBC assays

The MIC/MBC of rifampin, rifapentine, and rifabutin for *M. tuberculosis* H37Rv was determined by the broth macrodilution method. Frozen bacterial stocks were thawed and cultured in liquid growth medium until an optical density at 600 nm (OD_600_) of 0.8-1 was achieved. Cultures were then diluted 10-fold in assay media such that the desired assay inoculum (around 5 log_10_ CFU/mL) would be achieved by adding 100 µL of this bacterial suspension to each assay tube. Drug-containing assay media (2.4 mL) was dispensed into 14-mL round-bottom polystyrene tubes. Two-fold concentrations of each drug were evaluated at the following concentration ranges: rifampin, 0.0625-4 µg/mL; rifapentine, 0.01563-1 µg/mL; rifabutin, 0.00098-0.5 µg/mL. The final DMSO concentration was 4% (v/v) in all samples (including the no drug control). After addition of the bacterial inoculum (100 µL), samples were vortexed and then incubated without agitation at 37°C for 14 days. Ten-fold dilutions of the bacterial inoculum were prepared in phosphate-buffered saline (PBS) and cultured on non-selective 7H11 agar (500 µL per agar plate). On Day 14, the visual MIC was defined as the lowest drug concentration that prevented visible growth as assessed by the unaided eye. For all samples in which no visible growth was evident on Day 14 (*i.e*., samples at concentrations ≥MIC), as well as for the sample at the concentration just lower than the visual MIC, ten-fold dilutions were prepared in PBS and cultured on non-selective 7H11 agar (500 µL per agar plate). CFUs were counted after agar plates were incubated for 3-4 weeks at 37°C. The MBC was defined as the lowest drug concentration that decreased bacterial counts ≥2 log_10_ CFU/mL (*i.e.*, 99%) relative to the starting inoculum.

### Mice

All procedures were approved by the Johns Hopkins University Animal Care and Use Committee. Female or male adult BALB/c mice were purchased from Charles River Laboratories. Uninfected mice used for PK studies were maintained in animal biosafety level (ABSL) -2 facilities, and infected mice used for PK/PD studies were maintained in ABSL-3 facilities. All mice were housed in individually ventilated cages (up to five mice per cage) with access to food and water *ad libitum* and with sterile shredded paper for bedding. Room temperature was maintained at 22-24°C, and a 12-h light/dark cycle was used. All mice were sacrificed by intentional isoflurane overdose by inhalation (drop method) followed by cervical dislocation.

### PK studies

Uninfected 8-10 week old female (n = 84) and male (n = 48) BALB/c mice were used for PK studies. Rifapentine was dosed at 3, 1, 0.3, and 0.1 mg/kg in female mice and at 3 and 0.3 mg/kg in male mice. Rifabutin was dosed at 3, 1, and 0.3 mg/kg in female mice and at 3 and 0.3 mg/kg in male mice. Rifabutin was not dosed at 0.1 mg/kg as we expected plasma exposures would be below the lower limit of quantification. Drugs were administered once daily with nine total doses administered (dosing Monday-Friday during the first week and Monday-Thursday during the second week). Each dosing group included 12 mice. After the first dose (Day 0) and the ninth dose (Day 10), blood sampling was done by in-life mandibular bleeding at 0 h (just prior to the daily dose, mice 1-3) and at the following time points after the daily dose: 0.5 h (mice 4-6), 1 h (mice 7-9), 2 h (mice 10-12), 4 h (mice 1-3), 6 h (mice 4-6), 9 h (mice 7-9), and 24 h (just prior to the next day’s dose, mice 10-12). Blood was collected into BD vacutainer plasma separation tubes with lithium heparin, and plasma was separated by spinning at 15,000 rcf for 10 minutes at room temperature. Plasma was transferred to 1.5 mL O-ring screw-cap tubes and stored at −80°C until analysis. Frozen samples were shipped to the Infectious Disease Pharmacokinetics Laboratory at the University of Florida College of Pharmacy, where drug concentrations were measured by liquid chromatography-tandem mass spectrometry. The lower limit of quantification for rifapentine and rifabutin was 0.10 and 0.05 µg/mL, respectively.

### PK/PD studies

Two PK/PD studies were conducted using the paucibacillary mouse model of TPT, which has been previously described (19, 20, 30, 35). In this model, female BALB/c mice (6-8 weeks old) are immunized by aerosol infection with *M. bovis* rBCG30. Six weeks later, mice are challenged with a low-dose aerosol infection of *M. tuberculosis* H37Rv, and treatment is started six weeks after the challenge infection. All aerosol infections were performed using a Glas-Col Full Size Inhalation Exposure system. The bacterial suspensions used for each infection were prepared as described in **Table S8**. For each infection run, 10 mL of the appropriate bacterial suspension was added to the nebulizer per the manufacturer’s instructions. For the first PK/PD study, the aerosol infections were performed as follows: immunization, one infection run with 120 mice; first *M. tuberculosis* challenge infection, one infection run with 115 mice; second *M. tuberculosis* challenge infection, one infection run with 120 mice (see study scheme in **Table S3**). For the second PK/PD study, the aerosol infections were performed as follows: immunization, two infection runs with 70 and 69 mice; *M. tuberculosis* challenge infection, two infection runs with 70 and 69 mice (see study scheme in **Table S6**).

In the first PK/PD study, the *M. tuberculosis* challenge infection implanted lower lung CFU counts than anticipated, and one mouse sacrificed the day after infection had no detectable *M. tuberculosis*. Thus, to ensure that all mice were infected with *M. tuberculosis*, the mice were subjected to a second *M. tuberculosis* challenge infection. For logistical reasons, treatment was initiated 7 weeks after the second challenge infection. Prior to the start of treatment, mice were randomized into one of the 10 regimens described in **Table 2**. Data from the PK studies in uninfected mice were used to design rifapentine and rifabutin regimens that maintained plasma exposures around pre-determined target concentrations. In all groups, treatment was administered 7 days per week, and for BID regimens, the two daily doses were given 12 ± 2 hours apart. Treatment was originally planned to be administered for 6 weeks, with CFU and PK time points after 3 and 6 weeks of treatment. However, the COVID-19-related shut-down of Johns Hopkins University necessitated ending the study 1.5 days after the Week 3 time point. The original and final schemes for this study, which ended up including 125 mice, are presented in **Table S3**. On Day 0 and at Week 3, PK sampling time points for rifapentine-treated mice were 2, 9, and 24 hours after the first daily dose; and the sampling time points for rifabutin-treated mice were 1, 9, and 24 hours after the first dose. The PK sampling for week 3 was conducted 4 days before the sacrifice time point. Blood sampling, processing, and drug measurements were performed as described above for the PK studies. Lung CFU data were determined as previously described (30, 35), and the plating strategy and the dilutions used to determine the log_10_ CFU/lung for each mouse are provided with the individual mouse data. All CFU/lung (*x*) data were log-transformed as log_10_ (*x* + 1) prior to analysis. Differences in lung CFU counts relative to the positive control regimen were determined using two-way ANOVA with Dunnett’s multiple comparisons test using GraphPad Prism software version 9.3.0. The lung CFU counts for rBCG30 are summarized in **Table S9**. All individual mouse CFU and PK data and the statistical analysis of CFUs are available in **Data File S2**.

The second PK/PD study followed the originally designed study scheme and included a total of 147 mice (**Table S6**). In this study, oral rifapentine and rifabutin regimens were designed to simulate the sharp rise and slow decline of drug exposures following an LAI dose. The regimens were designed to maintain trough plasma concentrations above pre-determined target concentrations for 4 weeks following each simulated LAI dose. To achieve these dynamic exposure profiles, the oral dose was changed every 4 days of treatment, as detailed in **Table S5**. These regimens were designed based on the cumulative PK data generated in this project and on the previously published modeling work for rifapentine (6). 1HP was used as the positive control regimen, with isoniazid (10 mg/kg) administered 1 hour after rifapentine (10 mg/kg). Mice were randomly assigned to a regimen prior to the start of treatment. In all groups, treatment was administered 7 days per week, and for BID regimens, the two daily doses were given 12 ± 2 hours apart. Mice were sacrificed to determine lung CFU counts after 2, 4, and 8 weeks of treatment. CFUs were determined as described for the first PK/PD study, and at Week 8, lung homogenates were also plated on selective 7H11 agar containing 1 μg/mL rifampin to assess rifamycin susceptibility at the end of treatment. The plating strategy and the dilutions used to determine the log_10_ CFU/lung for each mouse are provided with the individual mouse data. Differences in lung CFU counts across regimens at each time point were determined using two-way ANOVA, full model with interaction term and Tukey’s multiple comparison test using GraphPad Prism software version 9.3.0. In mice that received a simulated LAI regimen, PK sampling was performed at Weeks 2, 4, 6, and 8 of dosing. Sampling was done 4 days before the sacrifice time point. In mice that received regimens with two simulated LAI doses, there was an additional sampling time point at Week 4 + 1 day, to capture the increased exposures associated with the simulated second LAI dose. Blood was collected, processed, and analyzed as described for the first PK/PD study. The lung CFU counts for rBCG30 are summarized in **Table S10**. All individual mouse CFU and PK data and the statistical analysis of CFUs are available in **Data File S3**.

### PK and PK/PD analyses

PK and PK/PD data analyses and simulations were carried out in the R programming environment (v 4.0.3) (36). Fitting of models to observed data made use of the Pracma library (37), with parameter estimation by nonlinear regression using the “lsqnonlin” function for nonlinear least squares optimization, with an objective function weighted by 1/(predicted value)^2^. Data from all individual mice in any given dataset were treated as a naïve pool (38) rather than using an average value at a given time point. Both rifapentine and rifabutin plasma PK exposure data were adequately described with one-compartment PK disposition models with 1^st^ order absorption input, parameterized with apparent clearance (CL/F), apparent volume of distribution (V/F) and 1^st^ order absorption rate constant (Ka). For study 2, simulated-LAI-like PK profiles achieved via oral dosing over 8 weeks, the apparent time dependent effect of an increase in CL/F over the timecourse was accounted for in the model by a factor for fractional (reported as %) cumulative increase in CL/F per dosing day. This method has been used previously (39), with the factor parameter named here as “Ind_factor” due to autoinduction being the likely cause, and was implemented by the following equation in the structural PK model:

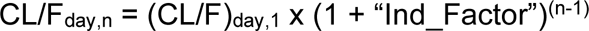

For timecourse profiles of bacterial load assessed by CFU count, the summary “net bacterial elimination” parameter “k_net_” (40) was estimated as the slope parameter of the modelled linear fitting though the log_10_ transform of the CFU profile. R code scripts used for analyses are available upon request.

## ACKNOWLEDGEMENTS

This research was funded by the National Institutes of Health (NIH) grant R61AI161809, and by a 2019 developmental grant from the Johns Hopkins University Center for AIDS Research, an NIH funded program (1P30AI094189), which is supported by the following NIH Co-Funding and Participating Institutes and Centers: NIAID, NCI, NICHD, NHLBI, NIDA, NIA, NIGMS, NIDDK, NIMHD. Pharmacokinetic modeling for rifapentine was supported by Unitaid project LONGEVITY (2020-38-LONGEVITY). The content is solely the responsibility of the authors and does not necessarily represent the official views of the funders.

